# Eugenol loaded lipid nanoparticles derived hydrogels ameliorate psoriasis-like skin lesions by lowering oxidative stress and modulating inflammation

**DOI:** 10.1101/2024.06.17.599274

**Authors:** Roshan Keshari, Abhay Tharmatt, Mamatha M. Pillai, Deepak Chitkara, Prakriti Tayalia, Rinti Banerjee, Shamik Sen, Rohit Srivastava

**Affiliations:** Department of Biosciences and Bioengineering, Indian Institute of Technology (IIT) Bombay, Powai, Mumbai, India; Department of Pharmacy, Birla Institute of Technology and Science-Pilani (BITS-Pilani), Vidya Vihar, Pilani, Rajasthan - 333031, India

**Keywords:** Eugenol, Soya lecithin, Psoriasis, lipid nanoparticles, keratinocyte, Imiquimod

## Abstract

Psoriasis is a chronic T-cell-mediated autoimmune skin disorder characterized by excessive epidermal thickening, keratinocyte over-proliferation, disruption of epidermal cell differentiation, and increased blood vessel growth in the dermal layer. Despite the common use of corticosteroids in psoriasis treatment, their limited efficacy and numerous side effects pose significant challenges. This research introduces a promising alternative approach by presenting hydrogels loaded with Eugenol (EU) in combination with Carbopol 974P (EUNPGel) for potential psoriasis management. EUN-loaded lipid nanoparticles (EUNPs) exhibit superior drug loading, enhanced release kinetics, long-term stability, and the ability to scavenge reactive oxygen species (ROS). Furthermore, EUNPs have been shown to inhibit keratinocyte proliferation, induce apoptosis, and augment the uptake of IL-6-mediated inflammation in human keratinocyte cells. Application of EUNPs-loaded gels (EUNPGel) to imiquimod-induced psoriatic lesions has demonstrated effective dermal penetration, suppressing keratinocyte hyperplasia and restoring epidermal growth. This led to a remarkable reduction in the Psoriasis Area and Severity Index (PASI) score from 3. 75 to 0. 5 within five days. These findings highlight the potential of EUNPGel as an innovative nanomedicine for treating inflammation. This novel approach enhances ROS scavenging capacity, improves cellular uptake, facilitates skin penetration and retention, reduces the activity of hyperactive immune cells, and suggests potential applications for treating other immune-related disorders such as acne and atopic dermatitis.

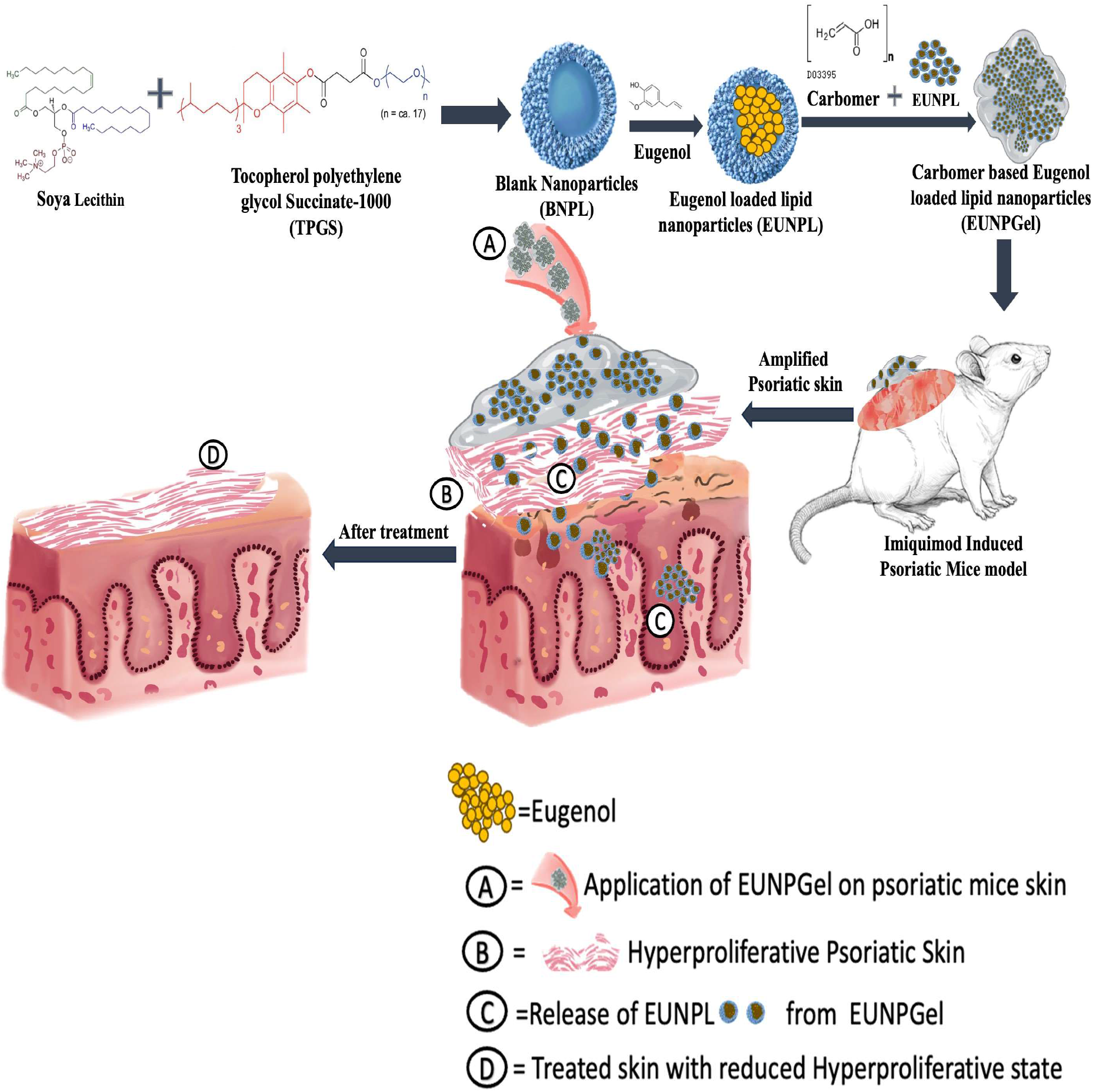

**Highlights:**

1. Hydrogel loaded with eugenol is an innovative alternative for psoriasis management.
2. Superior drug loading, release kinetics, stability, and ROS scavenging capacity.
3. Curb (human keratinocyte) HaCaT cells proliferation, induce apoptosis, lower IL-6 mediated inflammation.
4. Effective dermal penetration and retention both in vivo and ex vivo.

## 1. Introduction

Psoriasis is a recurring T-cell-mediated immune skin disorder whose prevalence drastically increases globally^1,2^. It occurs due to dermal acanthosis, proliferative nature of keratinocytes, alteration of epidermal differentiation, dermal angiogenesis and dense inflammatory cells on the dermal and epidermal layer ^3^. In standard body mechanisms, keratinocytes migrate to the epidermal layer within 4-5 days, and the exact mechanism happens 10-fold faster in psoriatic individuals, leading ^4^ to the development of inconsistent structure with parakeratosis by stratum corneum. However, the exact cause and mechanism are still unclear. It was hypothesized that stimulating damaged or flowing immunocytes such as T cells, neutrophils, dendritic cells, and macrophages causes epidermis hyperproliferation, thickening, and angiogenesis ^5,6^. The symptoms are repeated scaly, itchy, painful plaque, which leads to skin cracking, bleeding and distortion ^7^. Unfortunately, psoriasis can be controlled but not permanently cured ^8^. Currently, topical, systemic, biological agents and phototherapy are mainly used to inhibit the triggered dermal dendritic cells, which release IL-12 and IL-23 and further inhibit differentiating the T helper cells Th1 and Th 17 cells. These treatment regimens are mainly selected based on severity. However, among the various available therapies, topical treatments are widely accepted as standard care of psoriatic management because of low systemic toxicity and potential targeting ability.

Eugenol (1-allyl-4-hydroxy-3-methoxybenzene) is the primary ingredient of Ocimum gratissimum, a phenolic compound ^9^. It has a broad therapeutic activity such as analgesic ^10^, anti-inflammatory ^11^, anti-bacterial ^12^, anti-fungal ^13^, and anti-viral^14^. It is also reported that eugenol has antioxidant activity ^15^, and psoriatic patients often report redox imbalance, which implies the therapeutic potential of eugenol in psoriasis. These results led us to form the hypothesis that topically applied eugenol-encapsulated in lipid nanoparticles would penetrate the disrupted skin layer, internalize inside the hyperactivated keratinocytes, break the cycle of hyperproliferation of cells, and further improve the phenotypic psoriatic indexes. Additionally, it is attractive to employ more effective drug delivery systems to tailor the drug carriers to enhance skin penetration and retention time while considering the acknowledged drawbacks of existing conventional treatment. To overcome the traditional issues, nanoparticles (drug carriers) are always ideal for transdermal distribution to the skin because they fluidize the stratum corneum regardless of size, shape, charge, and hydrophilic-lipophilic balance (HLB)^16^. It reduces the skin barrier system and allows easy drug skin permeation by accumulating and penetrating through the hair follicle, holding 0. 1% of the total skin surfaces^17^. Considering the challenges and requirements in the current treatment regime, lipid-based nanosystems are widely explored for psoriasis applications. Due to its lipidic content, it is considered less toxic compared to other nanocarrier. It offers various advantages like an encapsulation of both hydrophilic and hydrophobic drugs, stability, biodegradability, controlled release, ability to break the skin barrier system, and interaction with the lipids of stratum corneum, which makes them a promising choice for psoriasis ^18^.

Topical formulations, mainly corticosteroids, are the first line of treatment for mild to moderate psoriasis, but most have limitations in penetration and retention. These limitations include off-target, dose dumping, and burst release. To overcome these limitations, we developed eugenol-encapsulated lipid nanoparticles, which localized explicitly on the stratum corneum’s surface and allowed us to penetrate the drug to the deeper layers of the skin and significantly lessen skin damage and proliferation, which improved the recovery of psoriatic symptoms.

## 2. Materials and Method

### 2.1 Materials and cell lines

Soya lecithin (Soya phosphatidylcholine), dialysis membrane (12. 5 KDa), fetal bovine serum (FBS), antibiotic-antimycotic solution, 3-(4,5-dimethylthiazol-2-yl) −2,5-diphenyl tetrazolium bromide (MTT) reagent, Dulbecco’s modified eagle medium (DMEM), dimethyl sulfoxide (DMSO) and phosphate buffer saline (PBS) pH 7. 4 were purchased from HImedia laboratories; Tocopherol polyethylene glycol 1000 succinate (TPGS) was procured from matrix science private limited; eugenol, triethanolamine, sodium benzoate and carbopol 974P were purchased from Sigma Aldrich. AddexBio supplied human keratinocyte cells; 4′,6-diamidino-2-phenylindole (DAPI) (≥98%), Rhodamine 6G (>97%), and 2′-7′-dichlorofluorescein diacetatem(H_2_DCFDA) (≥97%) were purchased from Merck KGaA. IL-6 was obtained from R&D Biosystems. Imiquimod was purchased (5 % W/W; Glenmark Pharmaceuticals, India) from a local vendor. Every experiment used high-purity deionized water (resistivity 18. 2 M cm) purified by a Milli-Q Plus water purification system (Millipore, USA). keratinocyte cells were grown as an adherent monolayer at 37°C in 5% CO2 in DMEM enriched with 10% FBS and 1% antibiotic. After attaining a confluency of 70–80%, cells were subcultured.

### 2.2 Preparation of eugenol-loaded lipid nanoparticles (EUNPs)

A high-speed homogenization method was used to prepare eugenol-encapsulated lipid nanoparticles. Briefly, 0. 1 % sodium benzoate as a preservative was added in pre-heated 65°C– 70°C Milli-Q water and subjected to magnetic stirring at 700–800 RPM, and when it was solubilized, 2 % of SPC was added at the same temperature until it gave a translucent solution. 1% TPGS was added to the above-prepared solution under constant stirring for 15 minutes. When all the components were solubilized entirely, they were further transferred into the bath sonicator for 10 minutes to disintegrate into small particles. After completing this procedure, it was homogenized with a T25 homogenizer (T25 ULTRA-TURRAX®, IKA, Staufen, Germany) for 50 min by dividing it into five cycles within a gap of five minutes for each process. Again, 1.5 % eugenol was slowly added to the formulation during homogenization by jacketing with ice for 10 minutes to yield the final formulation, i.e. eugenol-loaded lipid nanoparticles (EUNPs).

### 2.3 Preparation of eugenol-loaded lipid nanoparticles into hydrogel system (EUNPGel)

Eugenol nanoparticle gel (EUNPGel) was synthesized using Carbopol 974P (0. 8% w/v). In Brief, Carbopol 974P was hydrated in eugenol nanoparticle solution (EUNPs) for 8 hours at a magnetic stirrer at 1000 RPM to get a uniform mixture. It was again stirred vigorously with a high-speed homogenizer until visual homogeneity was achieved, and the pH was adjusted to 6. 5 by using triethanolamine at a concentration of 6. 5 µL/ml. The morphology of the EUNPGel was determined using FEG-SEM to confirm its integrity after formulating it into the topical Gel.

### 2.4 Particle Characterization and Optimization

The formulation was optimized based on the average size of particles and their distributions, encapsulation efficiency, zeta potential, SEM, TEM imaging, and FTIR spectra of freshly prepared nanoparticle solutions with lipid + drug, surfactant + drug, lipid + surfactant, and the mixture of lipid, drug, and surfactant.

### 2.5 Particle size distribution

The Z-average, polydispersity index (PDI) and zeta potential of EUNPs were estimated by a dynamic light scattering (DLS) method using a Malvern Zetasizer Nano (Malvern Instrument Ltd., UK). The correlated functional data from laser scattering were specifically fitted, and the Z-average size was determined as a value of intensity based on the average size. Before analysis, the formulation was 100-fold diluted in double-distilled water.

### 2.6 Imaging of EUNPs and EUNPGel by TEM and FEG-SEM

The morphological features of EUNPs were investigated using transmission electron microscopy (TEM). The copper grid (300 mesh) was covered with a drop of diluted EUNPs and dried at ambient temperature. A drop of phosphotungstic acid (2%) was added, dried for 30 minutes and examined using a 300 V-operating TEM (FEI Tecnai G2, F30). FEG SEM also analyzed the morphology of EUNPs via the drop-casting method by diluting the EUNPs to 5-fold with miliQ on aluminium foil and allowing it to air dry for 24 hours at room temperature by coating with iridium and examined under the JEOL JSM-7600F FEGSEM.

### 2.7 Entrapment Efficiency (EE) and Loading Percentage

An ultra-centrifugation technique was used to investigate the entrapment efficiency and loading percentage of EUNPs. EUNPs was centrifuged (Sigma 4K15, Refrigerated high-speed benchtop centrifuge) at 20,000 RPM for 2. 5 hours, and the supernatant was discarded; pellet was reconstituted in milli-Q water. The reconstituted formulation was further diluted ten times with methanol, and drug entrapment efficiency was estimated using HPLC by using the chromatographic condition mentioned in point 7. Percentage drug loading and EE were calculated using the below:-

> **EE-(Drug in nanoparticles/Initial drug added)*100**

> **Drug loading capacity- (weight of the drug in nanoparticles/weight of the nanoparticles)*100**

### 2.8 In vitro drug release

The release kinetics of eugenol from EUNPs, EUNPGel, and the free EU (EU) were analyzed using the dialysis bag method in PBS as a sink medium. The formulations were transferred to a 12. 5 kDa dialysis membrane previously pre-soaked and dipped at 37 °C while continuously stirring in a sink medium. The required sample volume was taken at various time intervals ranging from 0. 5, 1, 2, 4, 6, 8, 12, 24, and 48 hours, and the sink medium was replaced with an equivalent volume. The total percentage of eugenol present was determined by the HPLC method using chromatographic conditions in point 7.

### 2.9 Fourier transform infrared spectroscopy

FTIR spectra were captured using a Cary 630 (Agilent) spectrometer paired with the ATR module in the 4000 to 650 cm^−1^ wavenumber range. Before each acquisition, background spectra were collected, and 100% v/v ethanol was used to clean the ATR detector diamond. On the ATR crystal, 100 mg portions of the EUNPGel, EUNPs, and (blank nanoparticles) BNPs were deposited and air-dried for 30 to 45 seconds. Sixty-four scans per measurement spectra were collected at an 8^−1^ cm spectral resolution. The instrument parameters were Apodization Happ Genzel, phase correction Mertz, and set method gain (218).

### 2.10 Rheological study

The rheological behaviour of EUNPGel was analyzed using a rheometer (ARES G2 TA Instruments, MCR 702 Anton Paar). A parallel disc with a 25 mm diameter was selected, and a 0. 5 mm interval was maintained between the probe and disc. With an equilibration time of 9 s at 25°C and a shear rate of 10 s^−1^, the shaft was rotated for 3 min. At 20 time points, corresponding viscosities (cP) were recorded. The final product’s viscosity (cP) was calculated as the average of the previous three viscosity readings. Continuous shear stress was used to analyze the rheological behaviours of the gel and estimate the shear rate (1/s) as a function of shear stress (mPa). The shear rate was varied from 0. 1 to 100 s/s with 180 points, and the associated shear stress (mPa) was recorded at 37 °C ^19^.

### 2.11 Texture analysis

The prepared EUNPGel was examined using the TA2/1000 probe on the CT3-1000 Texture Analyzer (Brookfield Engineering Labs, Inc.) to investigate the textural features in compression mode. The detailed parameter used is mentioned in **(Table Supp. S1)**. Two cone-shaped probes, male and female, were employed, with the female probe connected to the sample holder’s base and used as a gel holder and the male probe attached to the load and used for insertion into the gel. The test proceeded by descending the male probe towards the female probe filled with a 5-gram gel at the pretest speed at a set distance target. The gel started to deform at a rate of 2. 0mm/s with a zero trigger load. The male probe pulled up from the female probe at the same speed initially used for penetration, and the result was recorded on TexturePro software ^20,21^.

### 2.12 Stability Studies

Stability studies of EUNPGel were assessed for 1, 2, 3, and 5 months at different storage conditions like 4 °C, room temperature, and accelerated conditions, particularly 40°C and 75 % RH. The samples were collected at regular intervals mentioned above, and their visual observations, such as syneresis, consistency, colour change, phase separation, grittiness, and drug content analysis, were studied. For drug content analysis, 100 mg of EUNPGel was resuspended in 900 µl of methanol and allowed to resuspend using a vortex and bath sonicator. The drug was then centrifuged at 10,000 RPM for 15 min, and the supernatant was analysed using the HPLC method using chromatographic conditions in point 7.

## 3. In vitro cell culture studies on human keratinocytes (HaCaT) cell lines

### 3.1 Cellular internalization study

In vitro, cellular internalization was studied using Rhodamine 6G, a red fluorescent dye. Rhodamine 6G was added to EUNPs and followed the same preparation method as mentioned above, and the amount of R6G encapsulated in EUNPs was measured at 548 nm using a microplate reader (Tecan Switzerland). Shortly, HaCaT cells were seeded in a 12-well plate on a sterile coverslip at 50,000 cells/well density. When cells reached nearly 70% confluency, the media was discarded, cells were washed and incubated with free@R6G and R6G@EUNPs for 4 hours at 37°C in 5 % CO_2_. After incubation, cells were washed and fixed with 4% paraformaldehyde (PFA) at 4°C for 15 minutes. After fixation, paraformaldehyde was discarded, washed and incubated with lectin for staining cell boundaries for 12 hours. After staining, lectin was discarded and washed with PBS twice. The nuclei were stained with DAPI at RT for 10 minutes. Finally, samples were transferred on a glass slide and imaged using CLSM (Carl Zeiss, LSM 780)at an excitation/emission of 525 / 548 nm respectively. To further investigate the uptake profile of nanoparticles in an inflammatory cell, cells were stimulated with IL-6 (25ng/ ml) for 24 hours before the initiation of experiments and followed the abovementioned steps.

### 3.2 In vitro cytotoxicity effect of EUNPs

The cytotoxicity effects of EUNPs, BNPs, and a free eugenol in DMSO (EU) were determined using an MTT assay. Briefly, HaCaT cells were seeded at a density of 10,000 cells/well in a 96-well plate for 24 hours at 37°C in 5% CO_2_. After 24 hours, cells were washed with PBS and incubated with the abovementioned formulations at a concentration range of 10 µg – 300 µg/ ml for 24 hours. Following treatment, the medium was removed, washed with PBS and MTT was added at a concentration of 0. 5 mg/ ml and incubated for 4 hours. After 4 hours, MTT was discarded, formazan crystals were dissolved in DMSO and absorbances were recorded at 570 nm using a microplate reader (Tecan Switzerland). The cytotoxicity effect of EUNPs, BNPs, and EU was studied by using the following formula:

> **Cytotoxicity = [(optical density of samples)/(optical density of control)] *100**

### 3.3 Colony formation assay

A colony formation assay was carried out to check the inhibitory effect of EUNPs on HaCaT cells. Briefly, HaCaT cells were seeded at a low density, i.e. 5000 cells/well in 6 well plates and incubated for 24 hours at 37 °C in 5 % CO_2_ ^22^. After 24 hours, cells were incubated with EUNPs, BNPs, and EU. Every alternative day, media was discarded, cells were treated with the abovementioned groups until the cells grew to their visible colonies. When colonies were visible, cells were washed, fixed with PFA at 4°C for 15 min. After 15 min, PFA was discarded, washed, colonies were stained with 0.5 % crystal violet solution for 10 minutes and excessive stains were removed by washing thrice in running water and imaged. The content of the cells was dissolved in 200 µl of methanol and absorbance was taken at 555 nm using a multiscan sky plate reader ^23^.

### 3.4 Apoptosis Assay

HaCaT cells were seeded at a density of 30,0000 cells/wells in a 6-well plate and allowed to adhere for 24 hours at 37 °C in 5 % CO_2_. After 24 hours, the media was discarded, cells were incubated with EUNPs, BNPs, and EU. After treatment, cells were washed, trypsinized, centrifuged and reconstituted in 1X staining solution in binding buffer. Annexin V FITC conjugate with propidium iodide were used to quantify cellular apoptosis according to the manufacturer protocol (Cayman multiparameter apoptosis assay kit USA). The apoptotic cells were measured using a flow cytometer (BD Biosciences) and data were interpreted using FlowJo software.

### 3.5 Mitochondrial Membrane Potential Assay

Mitochondrial membrane potential (ΔΨ) was investigated by tetramethyl rhodamine ethyl ester (TMRE) staining to understand the effect of EUNPs on the polarisation of the mitochondrial membrane. Shortly, HaCaT cells were seeded at a density of 50,000 cells/wells in a 24-well plate and allowed to adhere for 24 hours at 37 °C in 5 % CO_2_. After 24 hours, media was discarded, cells were treated for 12 hours with EUNPs, BNPs, and EU before being exposed for 1 hour with 75 µM tert-butyl hydroperoxide. Cells were eventually tagged with TMRE and Hoechst following the manufacturer’s instructions (Cayman multiparameter apoptosis assay kit USA) and imaged with confocal microscopy (Carl Zeiss, LSM 780) at an excitation / emission of 540/595 nm respectively and their quantification was performed with Image J software.

## 4. Antioxidant and free radical scavenging activity: *In vitro and ex-vivo*

### 4.1 2,2-diphenyl-1-picrylhydrazyl (DPPH) *in vitro* antioxidant assay

The antioxidant and free radical scavenging activities of the EUNPs were analysed by the DPPH Assay method ^24^.A DPPH solution of 0. 1 mM in 95% ethanol was prepared and 3. 9 ml of DPPH solution and 100µl of EUNPs were mixed. Absorbance at 517 nm was recorded at 10 min, and free radicals antioxidant activities were analysed using the following formula:

> **Free radical scavenging activity = (1-(I-II)/III) *100**

### 4.2 Intracellular ROS determination

HaCaT cells were seeded at a density of 50,0000 cells/wells in a 12-well plate and allowed to adhere for 24 hours at 37 °C in 5 % CO_2_. After 24 hours, media was discarded and cells were pre-treated with EUNPs, BNPs, and EU for 6 hours and additionally 1 hour incubation with tert-butyl hydroperoxide (75 µM). Eventually, cells were washed, labelled with 25 µM DCFH-DA and incubated at 37 °C for 30 min in the dark. After 30 min, DCFH-DA was discarded, washed, stained with DAPI for 5 min and imaged with confocal microscopy (Carl Zeiss, LSM 780) at an excitation / emission of 485 / 535 nm, and their quantification was performed with Image J software.

### 4.3 Ex vivo ROS (Reactive Oxygen Species) study in psoriatic mice model

Psoriatic skin specimens from different treatment groups were stored at 4% formaldehyde. During the study, skins were embedded in OCT solution (Thermo Fisher Scientific, Waltham, USA), cryosectioned at 5-7mm thick, immobilized on silane-coated slides (BioMarq, India) and stored at 4°C. For the ROS study, sections were washed with PBS and incubated with 0. 5 mM DCFH-DA for 30 min at 37°C in the dark, restained with DAPI (5 µg/ ml) for 10 minutes, washed with PBS and visualized with an EVOS M7000 imaging system (Thermo Fischer Scientific) under 10x magnification and the mean fluorescence intensity was analysed with Image J software.

## 5. Ex vivo studies

### 5.1 Ex vivo skin penetration and deposition study

To measure the penetrability of EUNPGel and free eugenol gel (EUgel) in the dermis and epidermis, an ex vivo skin penetration study was carried out using skin extracted from the dorsal side of Swiss albino mice after scarification ^25^. The collected skin tissue was separated from the subcutaneous adipose tissue and attached tightly between the donor and receiver compartments using clamps ^26^. The receiver compartment was filled with PBS, with the stratum corneum facing towards the donor compartment. In the donor compartment, EUNPGel and an EUgel equivalent to EUNPGel were added and allowed to penetrate for 24 hours at 37°C under stirring at 700–800 RPM.

After 24 hours, aliquots were taken from the receiver’s compartment and the skin sample was removed, washed with PBS and dried. The tape stripping method (cellophane tape locally purchased) was used to separate the stratum corneum. Briefly, the tape was attached to the skin so that the entire skin and tape were utilized; the first two pieces of stripped tape were discarded as they contained unabsorbed drugs. For separation of the stratum corneum layer, ten strips were detached and soaked in methanol for eugenol extraction for 24 hours at 4°C. The remaining skin was also minced into tiny parts and processed with a tissue homogenizer (T25 ULTRA-TURRAX®, IKA, Staufen, Germany) for 5 min at 5,000 RPM and transferred to the shaker at 4 °C for 24 hours ^27^. The sample was analysed by HPLC using the chromatographic condition mentioned in point 7.

### 5.2 Ex *vivo* imaging of psoriatic skin by IVIS system and their estimation by CLSM

The ex vivo distribution of R6G@EUNPGel and R6G@ EUgel in the psoriasis-prone skin of swiss albino mice was visualized using the rhodamine 6G (R6G) fluorescent dye. Briefly, R6G@EUNPGel was prepared as previously described above. Psoriatic mouse skin was positioned between the donor and receiver compartment of the Franz diffusion cells and firmly tightened with clips. The receiver chamber was filled with PBS and in the donor chamber, R6G@EUNPGel and R6G@ EUgel were placed, followed by incubation for 6, 12, and 24 hours at 37 °C with a stirring speed of 700–800 RPM. Skin samples were taken at each interval, carefully cleaned with PBS, and allowed to air dry. Cellophane tape was attached to the skin multiple times so that the entire skin and tape were utilized to remove the unabsorbed dye from the upper layer of the skin. The remaining tissue was imaged with an in vivo imaging system (IVIS) (Perkin Elmer UK) at an excitation / emission of 525 / 548 nm, respectively. To estimate the depth of nanoparticle penetration, vertical serial images of tissue were imaged at 5µm intervals, and X-Y orthogonal images (1024 x 132) were also taken using a Carl Zeiss, LSM 780 under 10x magnification and mean fluorescence intensity were analysed with Image J software.

## 6. In vivo studies on psoriatic mice model

### 6.1 Establishment of imiquimod-induced psoriasis-like mice model

All the experiments were initiated in strict accordance with the recommendation and approval of the Institutional Animal Ethics Committee (IAEC) (protocol no. IAEC/RES/31/10), and studies were carried out following CPCSEA guidelines. Before initiating the experiments, mice (8–12 weeks old) were acclimatized for one week under standard laboratory conditions. Imiquimod (IMQ), a ligand of Toll-like receptors (TLR), has been administered topically and has been shown to cause skin inflammation similar to psoriasis. On the experimental day, 62. 5 mg of imiquimod cream (5%) was topically applied to the skin of the right ear and the shaved area of the back for five days. During the model development stage, the mice were housed in separate cages to prevent them from scratching each other.

The therapeutic efficacy was analysed by the psoriasis area severity index (PASI), which is the gold standard approach for determining the degree of severity based on thickness, erythema and scaling ^28^. The values were noted by two independent colleagues on a scale of 0 to 4, where 0, 1, 2, 3, and 4 indicate nil, mild, moderate, severe, and very severe, respectively. The cumulative scoring, which includes erythema, scaling, and thickening, specifies the extent of inflammation on a scale from 0 to 12 ^29^.

### 6.2 In vivo therapeutic efficacy

Mice were randomly separated into five categories (n = 6); group one served as a negative control. Likewise, groups two to five received imiquimod topically to create psoriasis-like lesions. Briefly, group two was kept as a positive control without treatment; in groups three, four, and five, BNPGel, EUNPGel, and a EUgel of 100mg/cm^2^ of skin were topically applied. All the groups were treated continuously for five days, the efficacy of which was analysed daily using the psoriatic area severity index (PASI). Additionally, using a digital micrometre and a vernier calliper, respectively, the thickness of the right, left, and back skin was measured ^30,31^. Lastly, Mice were euthanized, back skin, right ear, spleen, and liver were isolated and stored at −80 for further experiments.

### 6.3 Spleen to body weight ratio

The spleen is considered one of the most significant immune-related organs. Its primary role is participating in immunological reactions, phagocytosis, elimination of old red blood cells, germs, or foreign antigens ^32^. An increase in the spleen’s weight relative to body weight indicates stimulation, proliferation and reflects the activation of immune cells in the spleen ^33^. At the end of the experiments, the mice spleens were isolated, weight and size were recorded quickly to avoid any errors due to dehydration.

### 6.4 Evaluation of *In vivo* cytotoxicity

To determine the in vivo cytotoxicity, the primary tissues of the different treatment groups such as liver, spleen, ear, and back skin were immediately dissected after euthanasia, and tissues were stored in 4% paraformaldehyde for further experiments. Changes in the Body weight of each mouse were monitored daily and the serum analysis levels of total bilirubin (T-BIL), alanine amino transferase (ALT), aspartate amino transferase (AST), blood urea nitrogen (BUN), and serum creatinine (SRE) were detected using a biochemistry analyzer (EM 200 Transasia).

### 6.5 Histopathological and immunohistochemical

Mice were euthanized, back skin, right ear, spleen, and liver were collected and stored in 4% paraformaldehyde for histological evaluation. Samples were dried in an ethanol gradient before being embedded in paraffin. Each tissue was sliced into 5-7 μm sections on glass slides, stained with hematoxylin / eosin and imaged using a bright field microscope. Specific parameters like angiogenesis, supra papillary thinning, inflammatory infiltrates, munro’s microabscess, a pustule of Kogoj, hyperkeratosis, parakeratosis, epidermal hyperplasia, inflammatory lesions, hyperkeratosis, parakeratosis, and total skin damage score (cumulative scoring generated after adding all the factors mentioned above) were investigated ^34^. Various liver parameters have been examined, like inflammatory cell infiltration, hepatocyte degradation, and overall liver damage ^35^. For spleen lymphocyte depopulation, splenocyte integrity and widespread spleen damage were also investigated.

For Immunohistochemistry analysis, skin specimens were deparaffinized, incubated with primary antibodies, including anti-Ki67,anti-NF-kB,anti-myeloperoxidase (MPO) and anti-CD4 and then again incubated with horseradish peroxidase (HRP) polymer conjugated anti-mouse and rabbit IgG and developed by 3,3. -Diaminobenzidine (DAB). Finally, randomly selected fields in each representative section of slide were chosen for immunohistochemical examination and examined under a light microscope.

## 7. Apparatus and chromatographic condition

An HPLC system made up of an isocratic pump (Jasco pump 4180, Japan), an autosampler (Jasco 4050, Japan), and a UV detector (Jasco MD, 4015, Japan) was used to analyse the samples. A sensitive, reliable method was developed with R^2^ of 0. 99 for all the experiments with the linearity range of 1-60 µg/ml. An Agilent ZORBAX SB-C18 column (4. 6 X 250 mm, 5um) was used for separation. The ratio of methanol to water in the mobile phase was 70:30 (v/v); injection volume 50 μl; flow rate 1. 0 ml/min; UV detection wavelength 286 nm. Data acquisition was performed using Chrom Nav chromatography data system software.

## 8. Statistical Analysis

The results of three independent experiments were expressed in terms of mean ± standard deviation or standard error of the mean (SEM). The statistical significance of the difference between different groups was determined via repeated measures of Analysis of variance (ANOVA) or student t-test analysis using GraphPad Prism software (version 8, San Diego, California, USA), where a p<0. 05 was estimated as statistically significant, p values are indicated by asterisks in figures (**p <0. 01,***p <0. 001****p <0. 0001)

## 9. Results & Discussions

### 9.1 Eugenol NP (EUNPs) loaded gels (EUNPGel) possess superior material properties, sustained drug release potential and long-term stability

For preparing Eugenol nanoparticles (EUNPs), different formulations were prepared by mixing core lipid (SPC), surfactant (TPGS) and eugenol in different ratios **(Table supp.2)**. After getting the desired size range of blank lipid NPs (200 − 300 nm, BNPs) ^36^, eugenol was incorporated via active loading through homogenization to obtain a homogeneous particle size of ≈ 200 nm and a PDI of 0.23 **(Fig. 1A)** with spherical in shape **(Fig. 1B)** having good colloidal stability (zeta potential −27 mV) **(Fig. 1C)**, and robust encapsulation efficiency (≈ 85%) and loading percentage (≈ 30.58 %). Consistent with colloidal stability, no aggregation was observed (**Fig. supp. 3**).

**Fig. 1:**
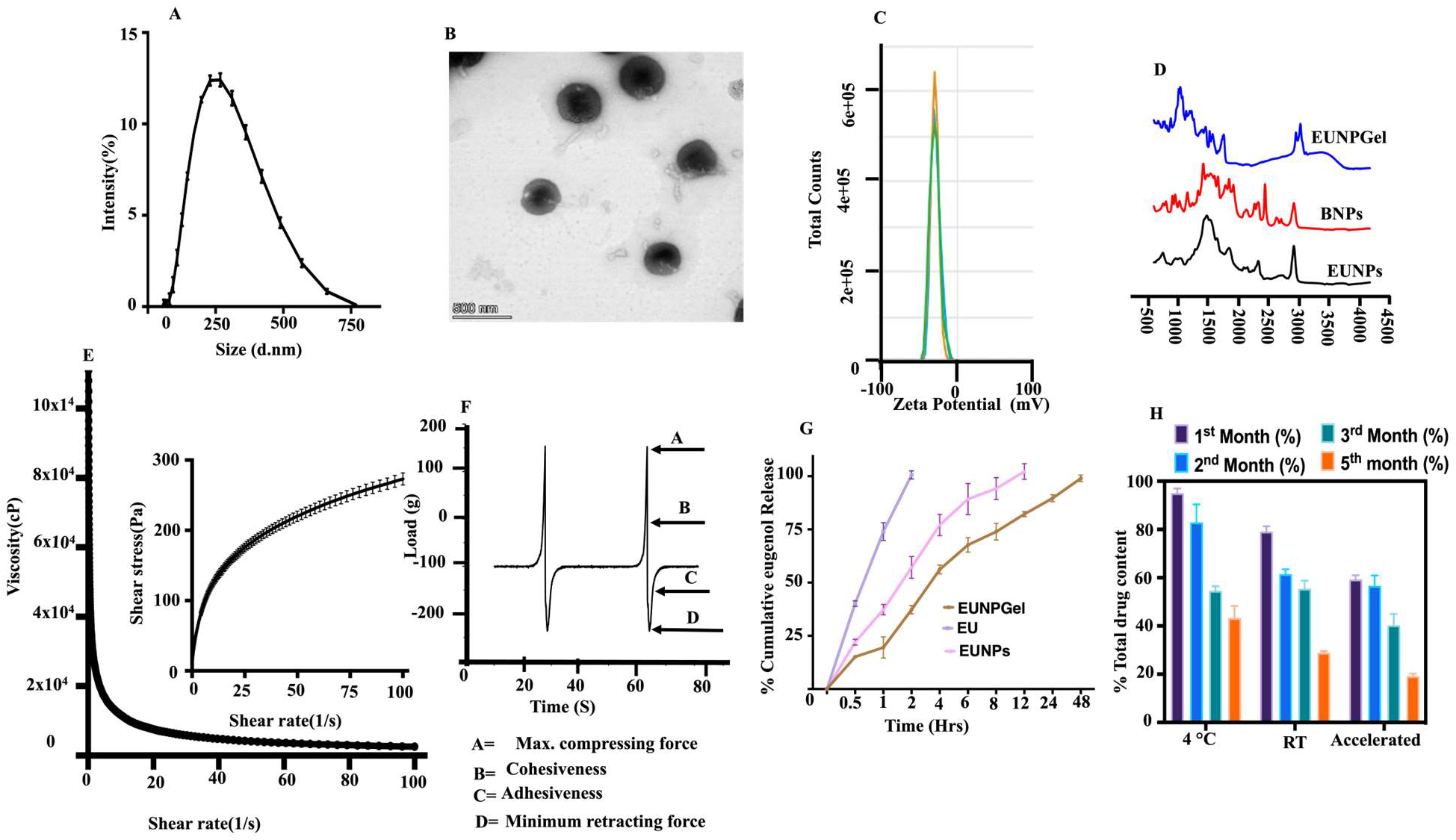
Physiochemical Characterisation of EUNPs, blank NPs (BNPs), EUNP loaded gels (EUNPGel) and free eugenol (EU). (A) Average hydrodynamic particle size distribution. (B) TEM image. (C) Zeta potential. (D) FTIR spectrum of EUNPGel, BNPs and EUNPs. (E) Representative plot of shear stress vs shear rate and viscosity versus shear rate graphs of EUNPGels. (F) Textural profile of the EUNPGel. (G) In vitro cumulative release profile of eugenol from EUNPGel, EUNPs, and EU solutions, (H) Stability study under various storage conditions. Data are represented as mean ± SD (n = 3)

EUNPGels were fabricated by combining EUNPs with carbomer overnight with constant stirring (see Methods for details). FTIR spectra of BNPs, EUNPs and EUNPGel revealed that there were no substantial structural changes as evident from the presence of both carboxylic acids of O-H stretching, alcohol of O-H stretching, and N-H stretching of amine salt along with C-H stretching of alkene and aldehyde **(Fig. 1D)**. EUNPGels exhibited shear thinning properties and varying thixotropic behaviour having viscosity of 12801 cp at 37 °C with a constant shear of 10s^−1^ **(Fig. 1E)** with texture analysis revealing good adhesivity and cohesiveness **(Fig. 1F, Supp. Table supp.4).**

In direct contrast to the burst release of eugenol from EUNPs and EU, sustained release of eugenol over 48 hours from EUNPGels may be attributed to passive adhesion or embedding of EUNPs to the tiny pores gel surface, thus making it difficult to diffuse and prevent the burst release of eugenol from EUNPGel **(Fig. 1G)**. The stability of EUNPGels assessed under various storage conditions up to 5 months revealed that the formulation was stable up to 2 months at 40 and room temperature (RT) but decreased to nearly half at the 3^rd^ and 5^th^ months **(Fig. 1H)**. Accelerated storage conditions (40°C, 75% RH) resulted in ≈ 50% reduction in the first month itself, possibly due to the volatile nature of eugenol. Syneresis, consistency, colour change, phase separation, and grittiness were also assessed. The colour changed to light brown at accelerated and RT storage conditions, possibly due to the degradation of SPC **(Fig. supp.5)**. Collectively, our results suggest EUNPGels possess robust mechanical properties, are stable and exhibit good release kinetics.

### 9.2 EUNPs induce cell death and inhibit proliferation of human keratinocyte cells (HaCaT cells)

Psoriasis is associated with increased proliferation of keratinocytes, inflammation and elevated ROS levels. Eugenol is a promising molecule for its anti-inflammatory, anti-oxidant and anti-proliferative properties^37^.To assess the internalisation efficiency of EUNPs by HaCaT cells, uptake experiments were performed with rhodamine (R6G) labelled EUNPs (R6G@EUNPs) and free R6G, respectively. R6G@EUNPs were readily internalized by keratinocytes as early as 4 hours, with 4 times faster uptake under IL-6 induced inflammatory conditions than healthy conditions **(Fig. 2B, C)**. MTT assay was performed by incubating cells with EUNPs, blank NPs (BNPs) and eugenol (EU) at 24 hours. It revealed the lowest cell numbers for the case of incubation with EUNPs **(Fig. 2D)**. These differences in cell numbers may be a consequence of increased cell death or decreased cell proliferation on EUNPs. Inhibition in colony formation **(Fig. 2E&F)** and the greatest fraction of apoptotic/necrotic cells in cells incubated with EUNPs **(Fig. 2G)** illustrates the pro-apoptotic and anti-proliferative effects of EUNPs on keratinocytes.

**Fig. 2:**
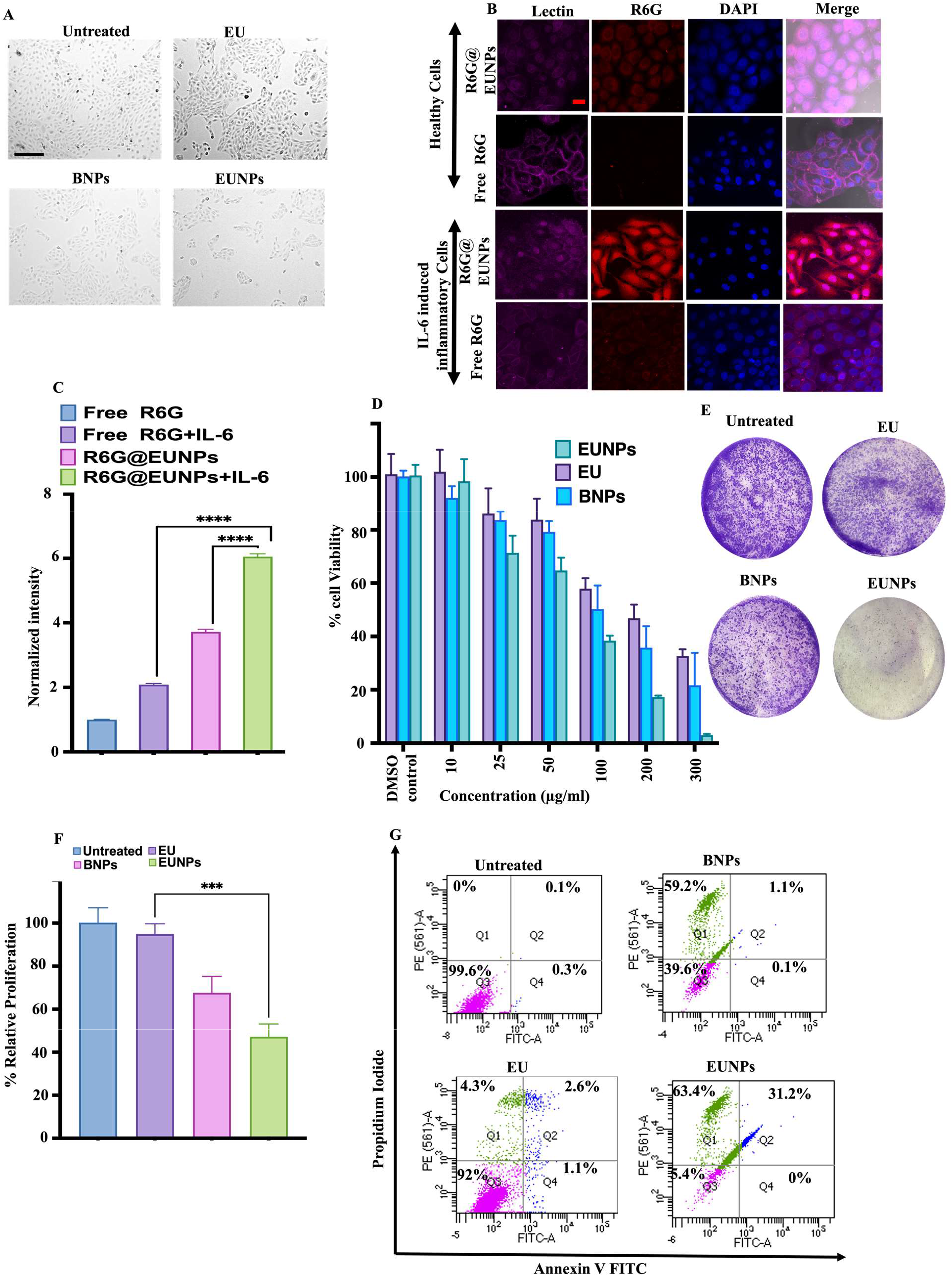
In vitro cellular assay on HaCaT cells. (A) HaCaT cells were imaged under the microscope after being subjected to EU, BNPs, and EUNPs (Magnification 20X, scale bar 275 µm). (B&C) Cellular uptake profile of R6G@EUNPs and free R6G in normal and diseased human keratinocyte cells and intensity quantification (Magnification 40X; scale bar 33µm). (D) cell viability assay. (E&F) Inhibition of cell proliferation by colony formation assay and their quantification. (G) Corresponding representative of FACS dot plot in late apoptotic cell (annexin V+/PI+) upon various treatments for 24 hours. Data are represented as mean ± SD (n = 3). P values were determined by using a t-test. ****p <0. 0001, ***p <0. 001

### 9.3 EUNPs suppress ROS and inhibit MMP activation

EU, a potent antioxidant ^38^, reduces persistent organic free radicals ^39^. By encapsulating EU in BNPs, their antioxidant properties were analysed by DPPH assay, which demonstrated robust ∼90% free radical scavenging activity of EUNPs. A ROS assay on keratinocyte cells was performed to further support this study using tert-butyl hydroperoxide (tBHP), a potent ROS generator. Earlier studies in vitro and in vivo models demonstrated that tBHP can damage keratinocytes through oxidative stress. Thus, using an exogenous stimulant of cellular oxidative stress, such as tBHP, can mimic a condition where skin cells are under increased oxidative stress ^40,41^. DCFH-DA fluorescence measurements were performed to quantify intracellular ROS levels. In contrast to high ROS levels observed in tBHP-treated cells, fluorescence was significantly decreased in cells treated with EUNPs, highlighting its anti-oxidant function **(Fig. 3A&B)**. ROS levels assessed ex vivo in excised skin of IMQ-induced psoriatic mice showed a similar trend **(Fig. supp. 6&7)**.

**Fig. 3.**
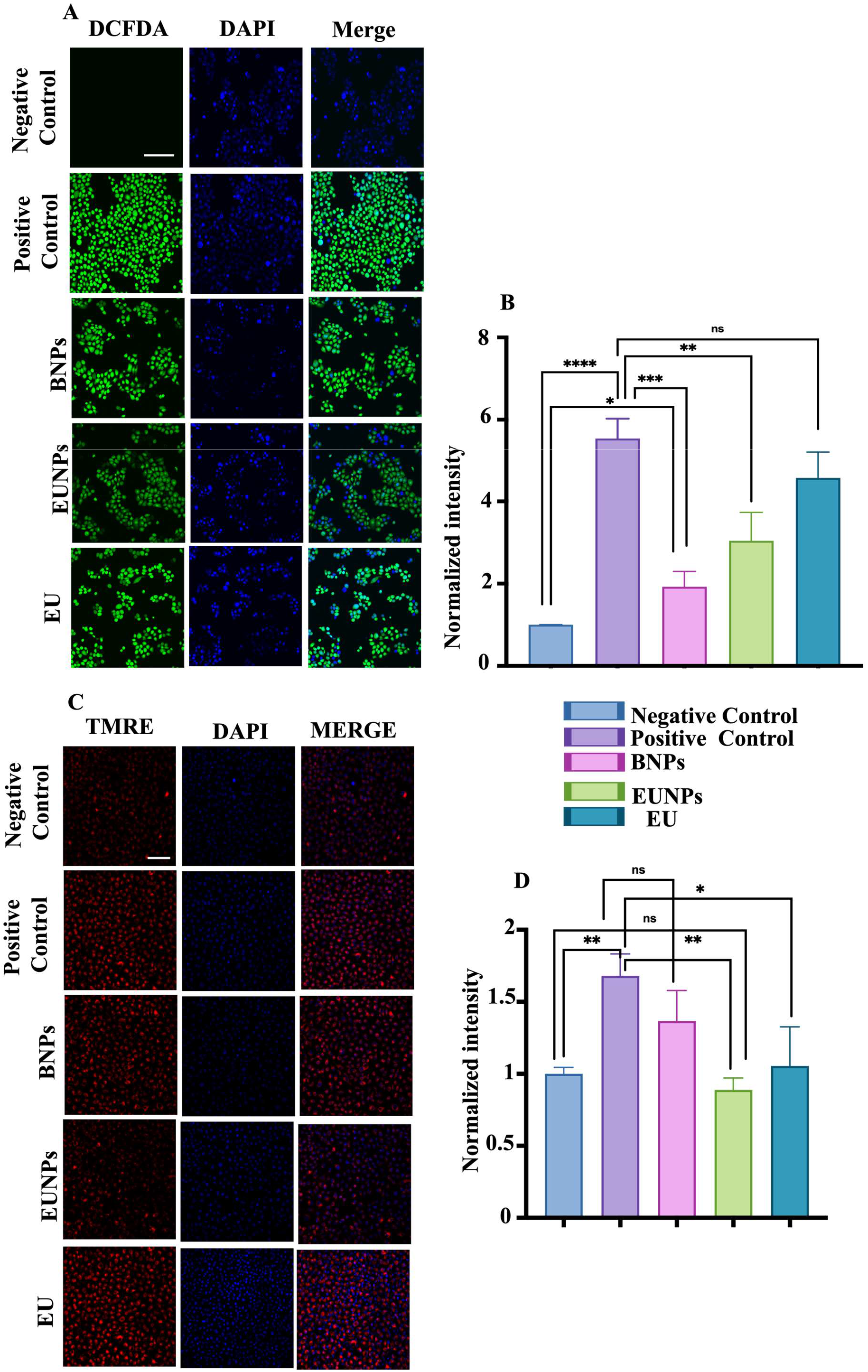
shows the Effects of EUNPs on tBHP-induced ROS generation and mitochondrial membrane potential depolarization. (A&B) Intracellular ROS level in different treatment groups and their intensity quantifications (Scale bar 132 µm).(C&D) TMRE staining indicating mitochondrial membrane potential and Quantification of intensity (Scale bar 104 µm). Data are represented as mean ± SD (n = 3). P values were determined by using a t-test. ****p <0. 0001, ***p <0. 001

Since increased ROS leads to mitochondrial damage ^44^, TMRE dye represents the changes in mitochondrial membrane potential and their alterations. The negative control showed red fluorescence, indicating standard mitochondrial potential. In contrast to high red fluorescence in positive control leading to mitochondrial depolarisation, fluorescence was significantly decreased in cells treated with EUNPs, restored as a negative control (**Fig. 3C&D)**, consistent with the ROS assay. Thus, the better performance in controlling ROS production and mitochondrial depolarisation by EUNPs might be due to the enhanced uptake of EUNPs.

### 9.4 EUNPGels can easily penetrate into deeper layers of psoriatic skin

To test the transdermal skin permeation capabilities of EUNPGel and EUGel, diffusion experiments were performed on excised psoriatic mice skin using the Franz diffusion apparatus **(Fig. 4A)**. Compared to EUGel, a 3-fold increase in drug accumulation in the receiver compartment was observed with EUNPGels indicative of increased drug penetration **(Fig. 4B)**. In line with this, 3 fold higher concentration of eugenol was detected in both dermis and epidermis in skin samples incubated with EUNPGels. In support of these observations, IVIS imaging revealed higher fluorescence of rhodamine (R6) at all time points in samples incubated with rhodamine loaded EUNPGels (R6G@EUNPGel) **(Fig. 4E**, S**upp. S8)**. Furthermore, confocal imaging revealed dye penetration to sub-cutaneous layers of the skin up to a depth of 80 *μ*m in EUNPGels, which was nearly double of that of free gel alone **(Fig. 4F, G, H&I)** (**Fig. supp. 9&10**). Collectively, these results illustrate the increased penetration ability of EUNPGels.

**Fig. 4:**
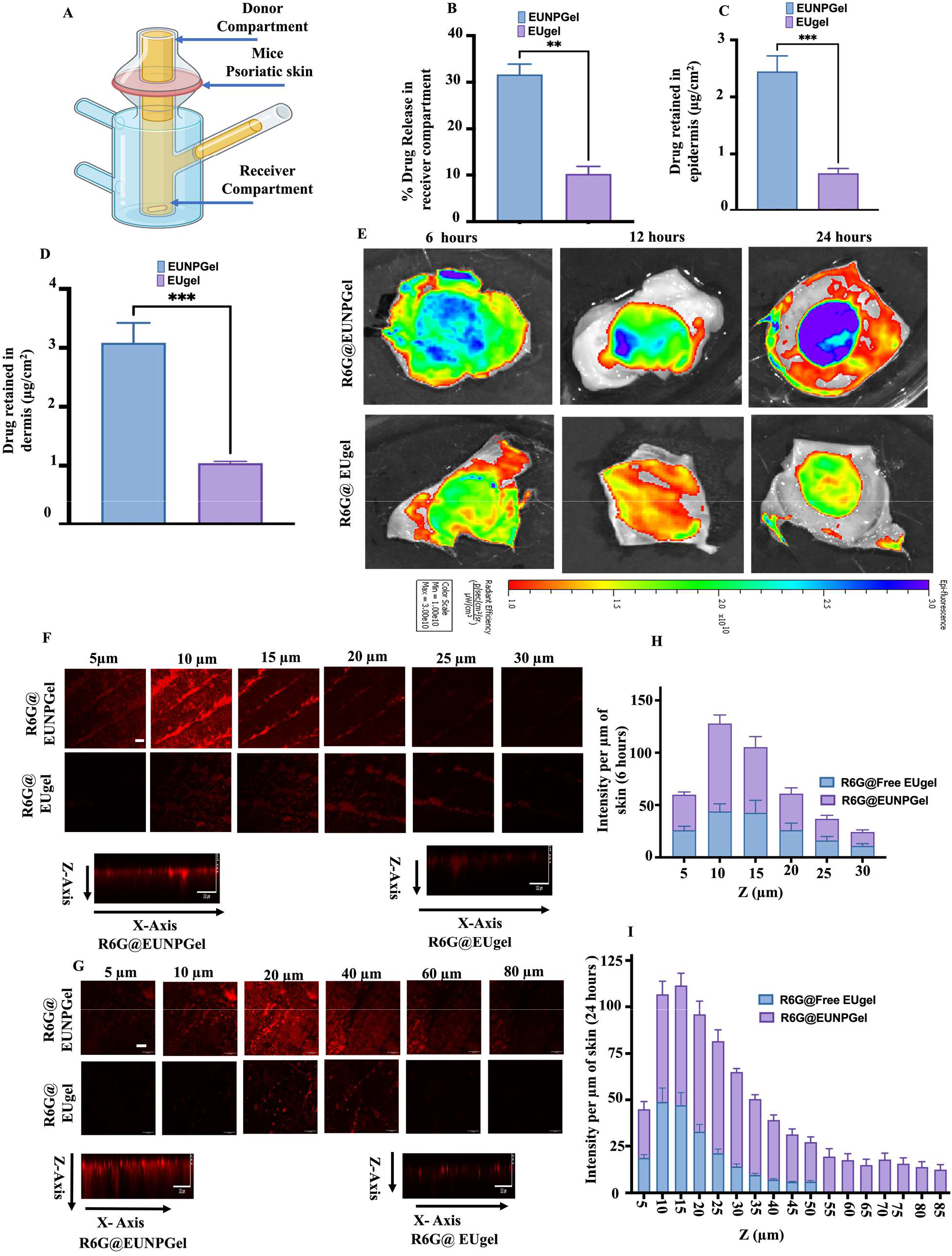
Ex vivo penetration, quantification and infiltration of drug and R6G dye in excised psoriatic mice skin and their bioimaging via IVIS system and confocal microscopy. (A) depicts the experimental setup with the Franz diffusion apparatus. (B, C&D) The amount of drug released in the receiver compartment and the amount of drug detected in the epidermis and dermis, respectively. (E) Ex vivo bioimaging of psoriatic skin treated R6G@ EUgel and R6G@EUNPGel for 6,12, and 24 hours. (F, G, H&I) representing serial confocal images of psoriatic skin tissues topically treated with R6G@EUNPGel and R6G@ EUgel for 6 and 24 hours and their intensity measurement, respectively. Images were obtained at 5 µm intervals on skin surfaces: scale bar, 132 µm. Data are represented as mean ± SD (n = 3). P values were determined using a t-test. **p <0. 01,***p <0. 001.

### 9.5 EUNPGels ameliorate psoriatic symptoms in imiquimod-induced mouse psoriasis model

Finally, we evaluated the therapeutic efficacy of EUNPGels using an imiquimod (IMQ)-induced psoriatic model, which suitably mimics skin abnormalities and signs of inflammation closely related to human psoriasis, including thickened epidermis, abnormal keratinocyte-related protein expression, inflammatory cell infiltration, and the secretion of proinflammatory cytokines ^45,46^. In this model, mice were treated with IMQ along with different formulations over a period of 5 days (**Fig. 5A, B)**. The degree of severity and therapeutic efficacy was analysed based on the PASI score. The topical administration of IMQ on mice back and ear skin for five consecutive days led to the formation of psoriatic-like lesions such as erythema, scaling and enhanced skin thickening **(Fig. 5C)**. It was noted that in the positive control, the significant psoriatic symptoms like scaling, erythema, and thickening started to increase from day 1 to 5 progressively, with the cumulative score crossing 10 at Day 5. Though both BNPL and EUGel treatment led to a decrease in disease severity, EUNPGel treatment led to the maximum therapeutic efficacy, with the PASI score reducing below 2. Consistent with this drop, epidermal tissue thickening—driven by inflammation and hyperproliferation of keratinocytes observed in mice given IMQ treatment only, was nearly abolished in mice treated with EUNPGels **(Fig. 5D, E)**. Also, loss in body weight observed in IMQ treated group was not observed in the other treatment groups **(Fig. 5F**). Increase in size and weight of spleen driven inflammation and accumulation of cells, was observed in IMQ treated mice, but abrogated to the highest extent in the group treated with EUNPGel to levels comparable to the negative control.

**Fig. 5:**
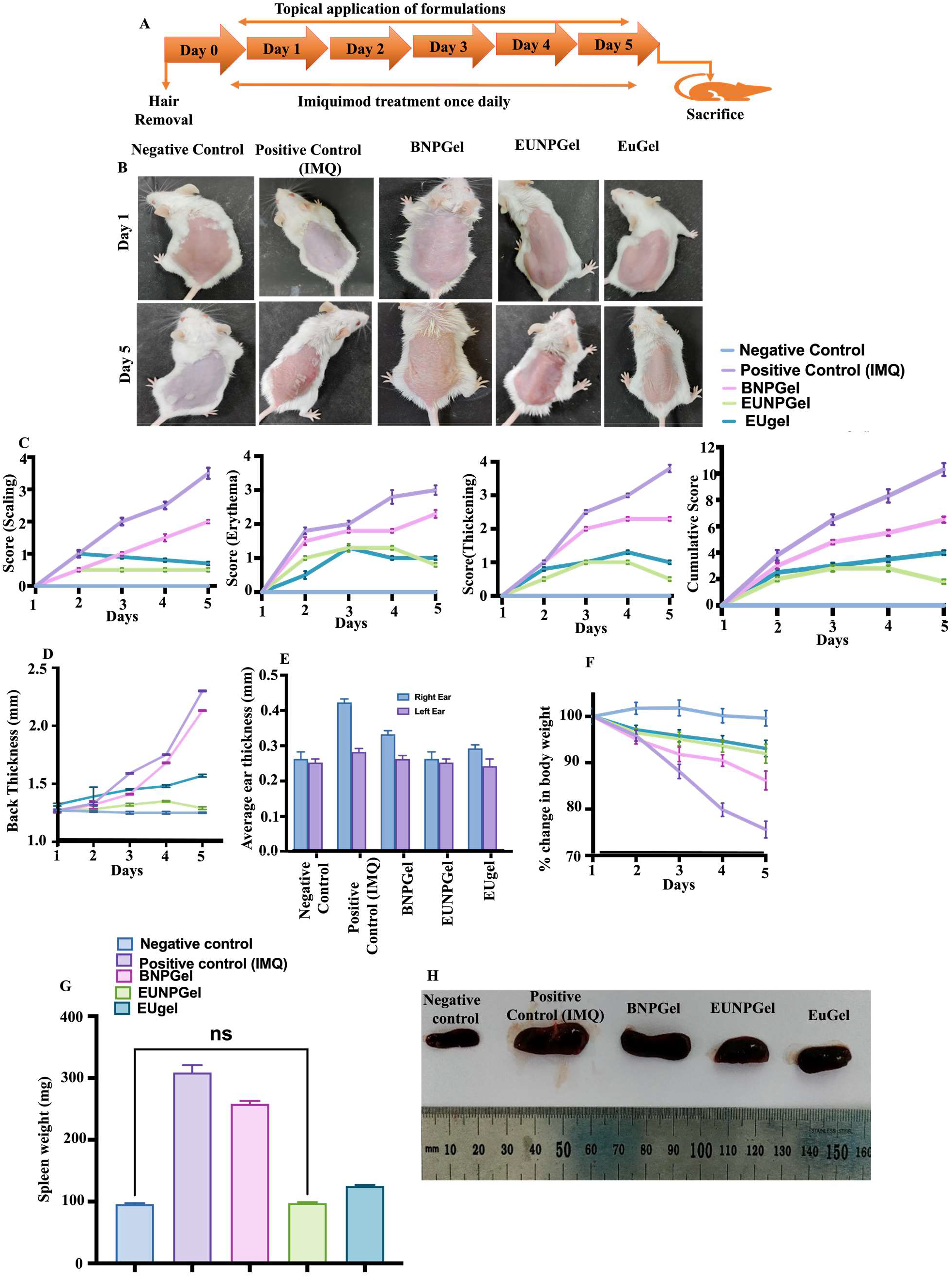
EUNPGel ameliorate (IMQ) induces psoriatic-like skin lesions in mice models. (A) Phenotypical representation of mice’s back skin, IMQ application, and treatment duration. (B)Clinical manifestation of each group on days 1 to 5. (C) scoring based on the clinical PASI score on a scale of 0–4 scaling, erythema, thickening, and total PASI score between (0–12). (D)Back thickness. (E) Comparison of thickness in the right and left ear. (F) Percentage body weight change. (G&H) Images of the spleen and the average weight of various treatment groups. Data are represented as mean ± SD (n = 3). P values were determined using a t-test. ****p <0. 0001, ***p <0. 001, **p <0. 01.

To understand how EUNPGel treatment reduced PASI score, histological analysis of the right ear, back skin, liver, and spleen were performed. The back skin rete ridges (red arrows) in IMQ treated mice indicate the induction of psoriasis-like lesions (Fig. 6A, B). However, mice treated with EUNPGel showed very minimal rete ridges, demonstrating the efficacy of EUNPGel in ameliorating psoriasis. Total skin damage was calculated by considering the major psoriatic skin parameters **(Tables 1&2),** which were minimal in EUNPGel with reduced epidermal thickness in the right ear **(Fig. 6C)** and back skin **(Fig. 6D)**, indicating the closest resemblance to normal skin. Similarly, histopathology of the spleen **(Fig. 6E)** and liver **(Fig. 6F)** were also studied, and total damage scores were noted, which were the lowest in the group treated with EUNPGel **(Tables 3&4).** Compared to other treatments, EUNPGel significantly improved the skin’s structure, reduced hyperproliferation and dorsal skin thickness and showed minimal liver and spleen damage.

**Fig. 6:**
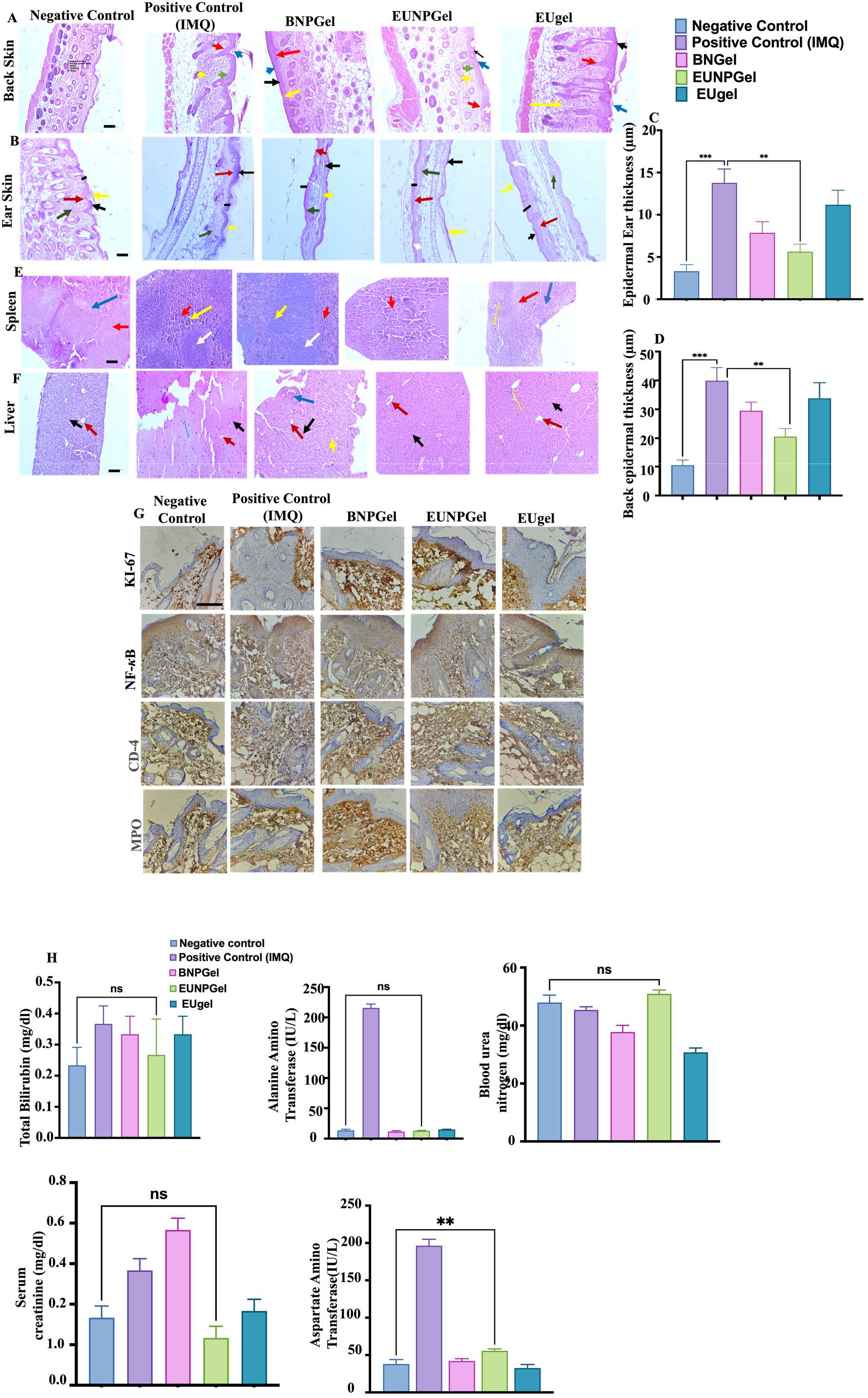
Histological, immunohistological, and in vivo cytotoxicity evaluation of mice tissue in each treatment group. (A, B, C, D) shows the H&E images of the back and ear skin and quantification of epidermal thickness. (E, F) shows the H&E images of the liver and Spleen.(G) Representative image for Ki-67, NF-κB, CD-4, and MPO immunohistochemistry in mice skin tissues (Scale bar 50 μm). (H) Evaluation of In vivo cytotoxicity via biochemical analysis of total Bilirubin (T-BIL), Alanine Amino Transferase (ALT), Blood Urea Nitrogen (BUN), Serum Creatinine (SRE), Aspartate Amino Transferase (AST) were measured in all treatment groups (Magnification: 20X). Data are represented as mean ± SD (n = 3). P values were determined using a t-test. ****p <0. 0001, ***p <0. 001, **p <0. 01. For interpretation of the references to colour in this figure legend, the reader is referred to the web version of this article). **Abbreviation** **Ear skin**: *Black arrow*; hyperkeratosis, *Yellow arrow*; parakeratosis, *Red arrow*; epidermal hyperplasia*, Green arrow*; infiltration of inflammatory cells, *White arrow*; microabscess. **Back skin**: *Black Arrow* ;hyperkeratosis, *Blue Arrow ;*parakeratosis, *Red arrow ;*epidermal hyperplasia, *Yellow arrow ;*inflammatory cells, *Green arrow ;*munro’s microabscess **Liver**: *Black Arrow ;*hepatocyte, *Red Arrow ;*central vein, *Yellow Arrow ;*degenerative changes, *Blue arrow* ;mononuclear infiltration **Spleen**: *Red Arrow*; red pulp, *Blue Arrow*; splenic vein, *White arrow*; white Pulp, Yellow arrow ; apoptotic Changes.

**Table-1.** Multiple signs reflecting the degree of skin injury following topical application of different treatment groups on the Back, Ear skin, spleen, and liver, respectively, of the imiquimod-induced psoriatic model. Data are represented as mean ± SD (n = 3)

| Groups | Hyperkeratosis | Parakeratosis | Epidermal Hyperplasia | Suprapapillary thinning | Inflammatory Infiltration | Microabscess | Pustule of Kogoj | Total Damage score |
| --- | --- | --- | --- | --- | --- | --- | --- | --- |
| Negative Control | 0 | 0 | 0 | 0 | 0 | 0 | 0 | 0 |
| Positive Control (IMQ) | ++ | ++ | +++ | +++ | +++ | ++ | + | 18 |
| BNPGel | + | + | ++ | ++ | ++ | + | 0 | 9 |
| EUNPGel | ++ | + | ++ | ++ | 0 | 0 | 0 | 7 |
| EuGel | ++ | ++ | +++ | +++ | +++ | ++ | + | 16 |

**Table-2.** Multiple signs reflecting the degree of skin injury following topical application of different treatment groups on the Back, Ear skin, spleen, and liver, respectively, of the imiquimod-induced psoriatic model. Data are represented as mean ± SD (n = 3)

| Groups | Hyperkeratosis | Parakeratosis | Capillary Proliferation | Epidermal Hyperplasia | Suprapapillary thinning | Inflammatory Infiltration | Microabscess | Pustule of Kogoj | Total Damage score |
| --- | --- | --- | --- | --- | --- | --- | --- | --- | --- |
| Negative Control | + | 0 | 0 | + | + | ++ | 0 | 0 | 05 |
| Positive Control (IMQ) | ++ | +++ | 0 | + | + | ++ | 0 | + | 10 |
| BNPGel | + | + | 0 | ++ | 0 | ++ | 0 | 0 | 06 |
| EUNPGel | 0 | 0 | 0 | + | + | + | 0 | 0 | 03 |
| EU Gel | + | 0 | 0 | +++ | ++ | + | + | 0 | 08 |

**Table-3.** Multiple signs reflecting the degree of skin injury following topical application of different treatment groups on the Back, Ear skin, spleen, and liver, respectively, of the imiquimod-induced psoriatic model. Data are represented as mean ± SD (n = 3)

| Groups | Mononuclear Infiltrations | Degeneration of Hepatocytes | Total Damage Score |
| --- | --- | --- | --- |
| Negative Control | 0 | 0 | 0 |
| Positive Control (IMQ) | + | +++ | 4 |
| BNPGel | + | ++ | 3 |
| EUNPGel | 0 | + | 1 |
| EUGel | + | + | 2 |

**Table-4.** Multiple signs reflecting the degree of skin injury following topical application of different treatment groups on the Back, Ear skin, spleen, and liver, respectively, of the imiquimod-induced psoriatic model. Data are represented as mean ± SD (n = 3)

| Groups | Depopulation of lymphocytes (Apoptosis Lymphocytes) | Degeneration of Red pulp | Total Damage score |
| --- | --- | --- | --- |
| Negative Control | 0 | 0 | 0 |
| Positive Control (IMQ) | ++ | + | 3 |
| BNPGel | ++ | 0 | 2 |
| EUNPGel | 0 | 0 | 0 |
| EUGel | + | ++ | 3 |

+ Indicate Mild damage ++ Indicate Mild moderate damage +++ Indicate Several damage

IHC analysis of critical markers, including Ki-67, MPO, CD-4 and NF-kB, correlated with the induction of psoriasis in IMQ-treated animals (Fig. 6G). Ki-67, a cell proliferation marker, has been proposed to identify psoriasis epidermal excessive proliferation ^48^. In tissue specimens, groups treated with IMQ showed a slight increase of 1-2% nuclear expression of Ki-67 in the epidermis’s acanthotic region, suggesting mild proliferation. However, the group receiving EUNPGel treatment demonstrated negative expression of Ki-67. Specifically in the NF-κB marker, the IMQ-treated group displayed robust NF-κB expression in the upper dermis inflammatory cells and non-specific cytoplasmic positivity. Still, the EUNPGel-treated group exhibited negative expression of NF-κB in both the epidermis and dermis, accompanied by non-specific cytoplasmic positivity. ^49^.

Similarly, in the CD-4 marker, groups subjected to IMQ only exhibited strong cytoplasmic expression of CD-4 in the inflammatory cells of the dermis, coupled with non-specific positivity in the extracellular matrix. EUNPGel displayed negative expression of CD-4 in both the epidermis and dermis, with non-specific positivity observed in the extracellular matrix. In the MPO marker, IMQ-treated groups exhibited cytoplasmic solid expression of MPO in the inflammatory cells of the dermis with negative expression in the epidermis. EUNPGel revealed negative expression of MPO in both the epidermis and dermis, coupled with non-specific cytoplasmic positivity. The group treated with EUNPGel demonstrated moderate cytoplasmic expression of MPO in the dermis along inflammatory cells with negative expression in the epidermis. ^50^.

Comparing these results with existing psoriasis literature, the observed changes in marker expression patterns influenced by EUNPGel treatment align with the modulation of critical pathways associated with psoriasis pathogenesis. These findings underscore the promising role of EUNPGel in addressing critical cellular and molecular aspects of psoriasis and call for further investigations to validate its potential clinical application ^51,52^.

### 9.6 In vivo cytotoxicity

In **(Fig. 5H)**, in vivo cytotoxicity demonstrated that the disease-induced mice model showed significant changes in liver enzymes like alanine aminotransferase, aspartate aminotransferase, serum creatine, total bilirubin and negligible effect in blood urea nitrogen in all the groups ^53^. However, groups treated with BNPGel and EUgel showed little changes in these enzymes, but a significant reduction was observed in the group treated with EUNPGel compared to the negative control group. The outcomes from these results indicate that topical application of EUNPGel substantially mitigates psoriasis-like inflammations to an extent.

## 10 Conclusions

These findings highlight the potential of eugenol as a novel nanomedicine for treating psoriatic-like inflammation. Loading of eugenol into SPC nanoparticles showed superior therapeutic efficacy, controlled release, enhanced stability, drug dispersion, reduced dose dumping and subsequently limited adverse effects as confirmed by in vitro, ex vivo, and in vivo. Similarly, In vitro studies on HaCaT cells, like colony formation assay, apoptotic assay, and enhanced cellular uptake in IL-6 mediated inflammations compared to healthy HaCaT cells, further supported the studies. An ex vivo study on psoriatic mice showed better drug absorption and deposition at various skin layers, which might be due to the occlusive effect of forming an intact thin film on the skin’s surface ^43^. However, this could also result from the EUNPGel accumulating skin appendages like hair follicles, which could serve as a long-term drug reservoir ^44^. Moreover, topical application of EUNPGel on an imiquimod-induced psoriasis model showed favourable effects on immune cells and keratinocytes and minimal systemic toxicity. Further, this can be expanded to deliver hydrophobic and hydrophilic drugs for treating other immune-related disorders such as acne and atopic dermatitis eczema and can be further translated for commercialization.

## 11 Authors Contributions

**RK-** Conceptualization, methodology, execution, investigation, analyzing experiments, literature survey, writing original draft, review & editing, **DC & AT-** *In* vivo study, and interpretation of histology data, **PT & MP -** Histology slide preparation and interpretation of IHC data, **Late RB-** Initial fund acquisition, supervision and conceptualization, **SS&RS**-Conceptualization, Supervision, Investigation, Project administration, Funding acquisition, Resources, review & editing of manuscript

## 12 Acknowledgement

The author acknowledges SAIF (Sophisticated Analytical Instrument Facility) and IRCC (Industrial Research & Consultancy Centre) at the Indian Institute of Technology Bombay, India, for providing instrumental support. Roshan Keshari also acknowledges the DAAD (German Student Exchange Programme), Govt. of Germany, for a research fellowship.

## 13 Declaration of competing interest

All authors declare no conflicts of interest.

## Notes

### Competing Interest Statement

The authors have declared no competing interest.

