## Supplementarty for "Eugenol loaded lipid nanoparticles derived hydrogels ameliorate psoriasis-like skin lesions by lowering oxidative stress and modulating inflammation"

**Table S1 -Test method used for the textural analysis of EUNPGel**

| **Parameters** | **Value** |
| --- | --- |
| Test Type: | TPA |
| Target: | 36.0 mm |
| Hold Time: | 0 s |
| Trigger Load: | 0 g |
| Test Speed: | 2.00 |
| Return Speed: | 2 mm/s |
| # of Cycles | 2.0 mm/s |
| Target Type | Distance |
| Recovery Time: | 0 |
| Same Trigger: | True |
| Pretest Speed: | 2.00 mm/s |
| Data Rate: | 30.00 points/sec |
| Probe: | TA2/1000 |
| Fixture: | TA-BT-KIT |
| Load Cell | 10000g |

**Table S2- Formulations and optimisation of EUNPL. Data are represented as mean ± SD (n = 3).**

| **Formulation Name** | **Size (nm)** | **Zeta (mv)** | **PDI** | **EE %** |
| --- | --- | --- | --- | --- |
| SPC+drug | 204.7±36.2 | 38.7±7.4 | 0.31 | 84.1±17 |
| TPGS+drug | 87.7±11.5 | 14.8±0.3 | 0.49 | 82.4±5.4 |
| SPC+TPGS | 353.3±29.6 | 35.9±0.9 | 0.24 | --- |
| Final formulation | 220.6±16.9 | 35.2±1.9 | 0.26 | 85.58±6.5 |

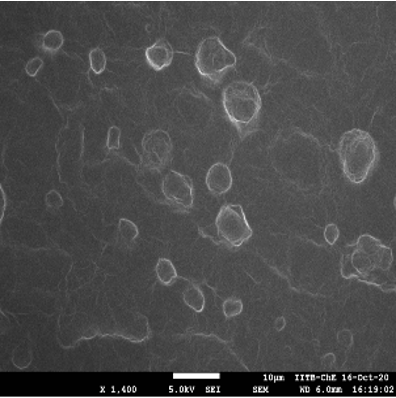

**Fig. S3: SEM image of EUNPL**

**Table S4. Texture Analysis profile EUNP Gel Data are represented as mean ± SD (n = 3)**

| **Hardness (g)** | 274. 00 ± 29. 50 |
| --- | --- |
| **Adhesive force (g)** | 133. 00 ± 15. 4 |
| **Adhesiveness (mJ)** | 4. 60 ± 0. 50 |

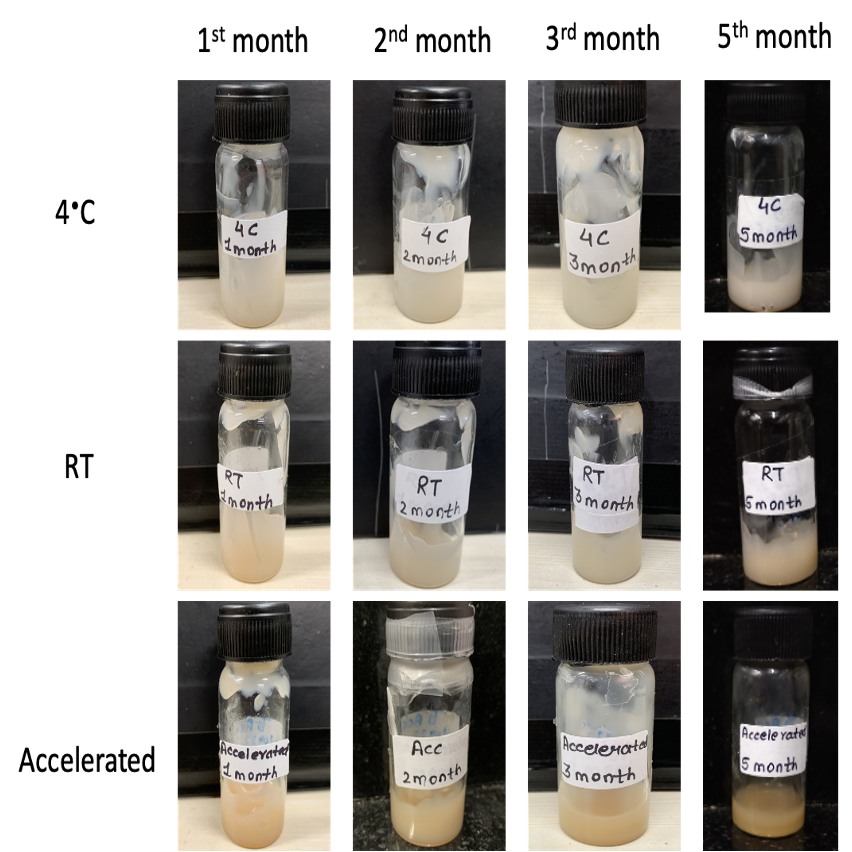

**Fig.S5: Photographic image of EUNPGel at different months and their conditions**

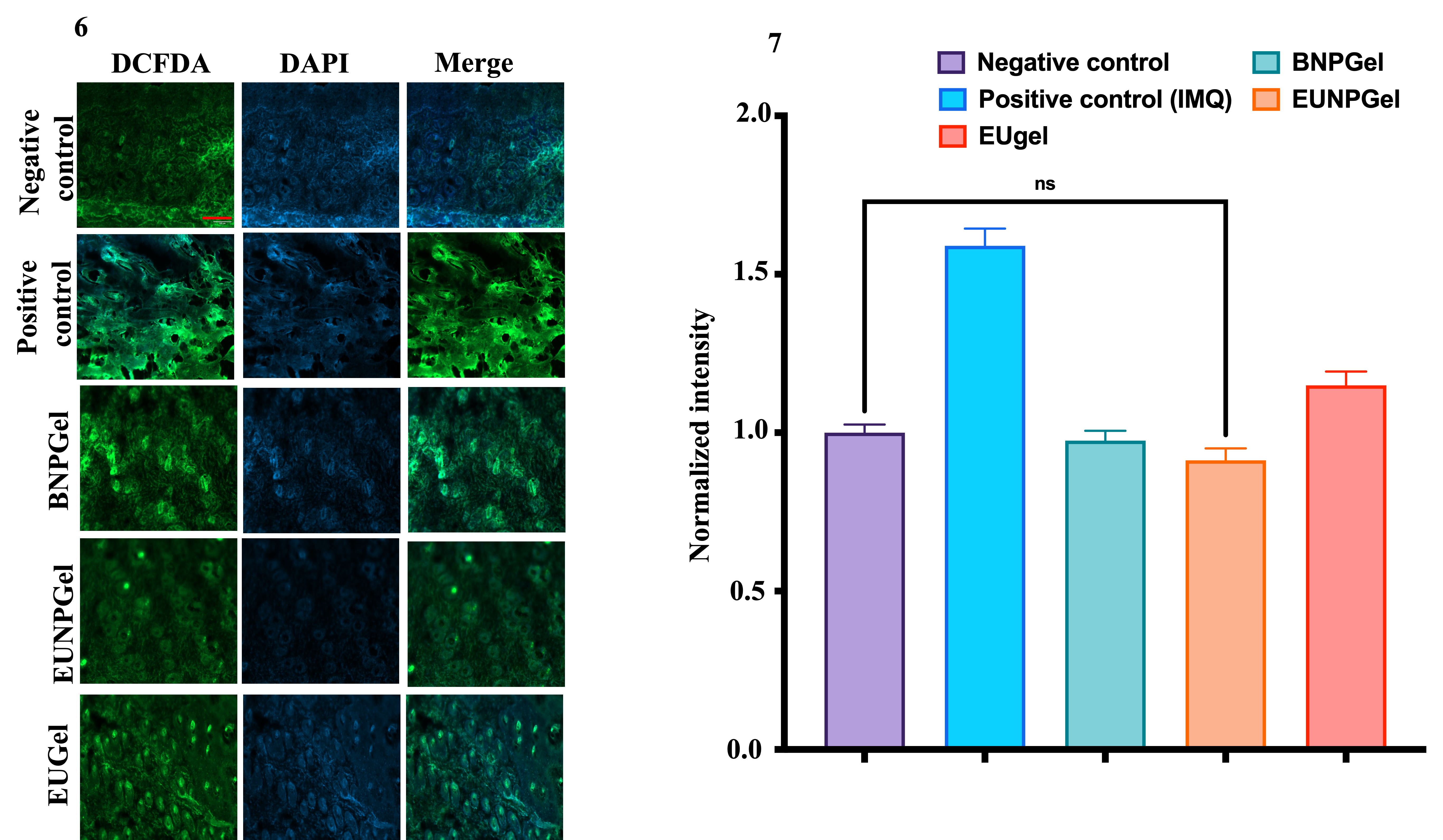

**Fig. S6&7: ROS study on excised skin of IMQ-induced psoriatic mice model and their quantification with Image J software. Data are represented as mean ± SD (n = 3). P values were determined using a t-test. ****p <0. 0001, ***p <0. 001, **p <0. 01.**

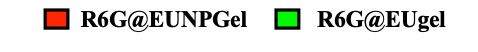

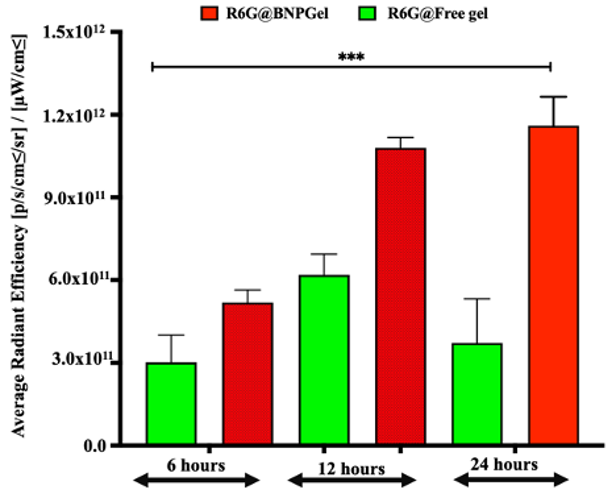

**Fig.S8: Quantification dye penetration at various intervals with Image J software. Data are represented as mean ± SD (n = 3). P values were determined using a t-test. ****p <0. 0001, ***p <0. 001, **p <0. 01.**

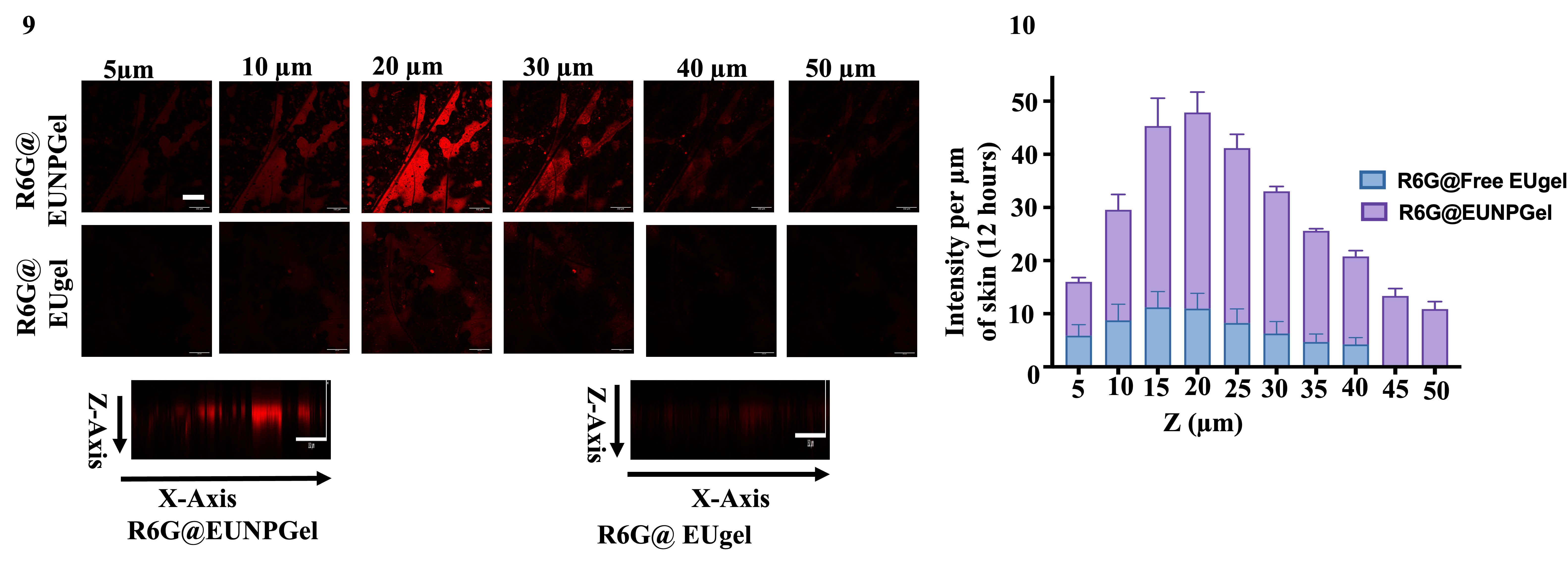

**(Fig.9 & 10) representing serial confocal images of psoriatic skin tissues topically treated with R6G@EUNPGel and R6G@ EUgel for 12 hours and their intensity measurement. Images were obtained at 5 µm intervals on skin surfaces: scale bar, 132** µ**m**. **Data are represented as mean ± SD (n = 3). P values were determined using a t-test. **p <0. 01,***p <0. 001.**
